# Enriched Methylomes of Low-input and Fragmented DNA Using Fragment Ligation EXclusive Methylation Sequencing (FLEXseq)

**DOI:** 10.1101/2024.11.28.625942

**Authors:** Jingru Yu, Lauren S. Ahmann, Yvette Y. Yao, Wei Gu

**Author notes:** Corresponding author: Wei Gu, Assistant Professor, Department of Pathology, Stanford University, 3373 Hillview Ave, room 220, Palo Alto, CA 94304-1204.

## Abstract

Methylome profiling is an emerging clinical tool for tumor classification and liquid biopsies. Here, we developed FLEXseq, a genome-wide methylation profiler that enriches and sequences the fragments of DNA flanking the CCGG motif. FLEXseq strongly correlates (Pearson’s r = 0.97) with whole genome bisulfite sequencing (WGBS) while enriching 18-fold. To demonstrate the broad applicability of FLEXseq, we verified its usage across cells, body fluids, and formalin-fixed paraffin-embedded (FFPE) tissues. DNA dilutions down to 250 pg decreased CpG coverage, but bias in methylation remained low (Pearson’s r ≥ 0.90) compared to a 10 ng input. FLEXseq offers a cost-efficient, base-pair resolution methylome with potential as a diagnostic tool for tissue and liquid biopsies.

---

Highly multiplexed DNA methylation testing is rapidly advancing, fueled by the growing interest in early cancer detection^1–4^ and the clinical adoption of tumor classification^5–8^. Further applications are emerging in diverse fields, such as aging through epigenetic clocks^9^ and liquid biopsies based on cell type deconvolution^10–12^. Methylation markers provide stable epigenetic data on DNA preserved in clinical specimens like formalin-fixed paraffin-embedded (FFPE) tissues and cell-free DNA (cfDNA) from body fluids.

Current methylation assays have limitations in clinical use. Whole-genome bisulfite sequencing (WGBS) is the gold standard but is not routinely used as it is cost-prohibitive. Reduced-representation bisulfite sequencing (RRBS)^13–16^ covers a fraction of the methylome and cell type-specific markers. Microarrays have high input requirements and lack linked data from single DNA fragments. Targeted methods require *a priori* knowledge of markers that rapidly evolve and differ across hundreds of unreferenced tumor types and backgrounds.

Here we present FLEXseq (Fragment Ligation EXclusive methylation sequencing), a methylation profiling method that targets the adjacent flanks of CCGG motifs in the genome that associate with cell type-specific markers, enhancers, and other epigenetic functional elements (Fig. 1). Extended-representation bisulfite sequencing (XRBS) also targets CCGG flanks^17^, but FLEXseq excels at fragmented and low-input DNA found in clinical specimens (e.g. cfDNA, FFPE tissue DNA). To design for fragmented DNA, we introduced a semi-permissive adapter that selectively blocks the free ends of non-target DNA, allowing FLEXseq to deliver a highly on-target and accurate methylome.

**Fig. 1:**
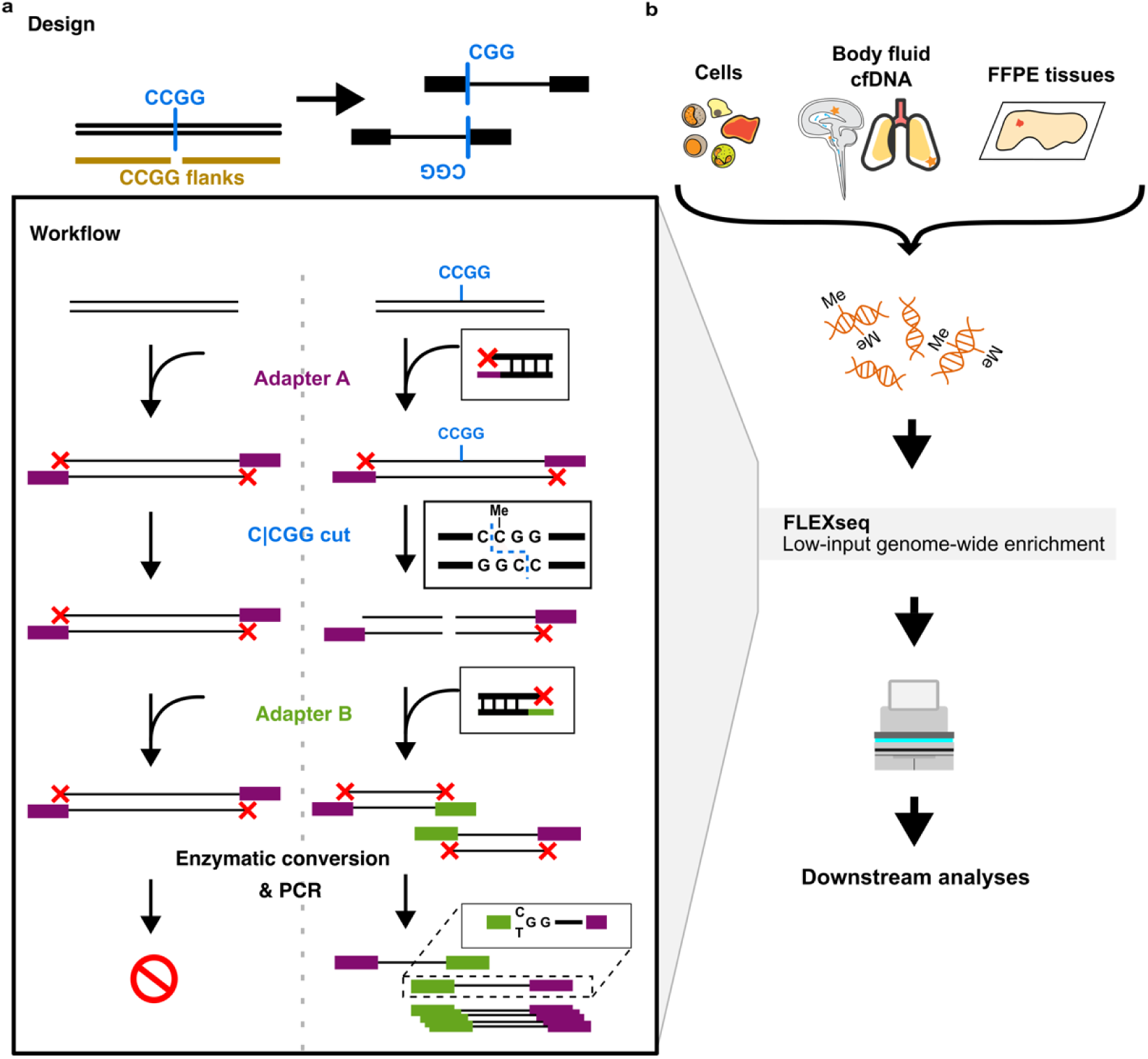
Schematic of FLEXseq design and workflow. The design goal and workflow of FLEXseq. The design aims to sequence adjacent regions flanking CCGG motifs while preserving the methylation markers at the motif. Information-poor regions are suppressed, while information-rich CCGG flanking regions are amplified. In the workflow, input DNA fragments are first ligated with a semi-permissive Adapter A, which serves both as an essential blocker for untargeted DNA and a required piece of targeted DNA. The nuclease MspI then cuts at the CCGG motif, followed by the ligation of Adapter B. Only molecules with both Adapters A and B are amplified and sequenced, excluding untargeted DNA with only one type of adapter.

## Results

### 1. Design and benchmarks of FLEXseq – enriched methylation profiling

#### FLEXseq - theoretical coverage

The need to detect and classify suspected tumors motivated us to design a genome-wide methylation profiler that overlaps with cell type-specific markers^11^ and references for tumor classification^6,8^. We started by calculating the theoretical overlap of various approaches with past references (Table 1), focusing specifically on short cfDNA (∼160 bp) and FFPE (∼100-500 bp) DNA in clinical specimens.

**Table 1.** *In silico* coverage of RRBS, XRBS, FLEXseq, methylation arrays, and MeDIP-seq.

| Method | MspI *in silico* digestion | Fragments | Coverage of CpGs | Coverage of filtered cell type-specific markers | Coverage of the top 100 cell type-specific markers | Coverage of CpGs in the top 100 cell type-specific markers |
| --- | --- | --- | --- | --- | --- | --- |
| RRBS | Maximum length 50 bp | 143,237 | 853,261 (3.3) | 16,905 (5.2) | 203 (5.3) | 844 (3.2) |
|  | Maximum length 100 bp | 260,753 | 1,967,891 (7.6) | 36,672 (11.4) | 448 (11.9) | 2155 (8.2) |
|  | Maximum length 150 bp | 319,553 | 2,960,586 (11.4) | 52,347 (16.3) | 629 (16.7) | 3566 (13.6) |
|  | Maximum length 200 bp | 363,551 | 3,762,719 (14.6) | 65,627 (20.4) | 786 (20.8) | 4680 (17.9) |
|  | Maximum length 300 bp | 391,103 | 4,790,769 (18.6) | 84,387 (26.3) | 976 (25.9) | 6064 (23.2) |
| CCGG Flanks | Extend 50 bp in each direction | 2,317,745 | 6,547,055 (25.4) | 149,321 (46.5) | 1,781 (47.2) | 8640 (33.1) |
|  | Extend 100 bp in each direction | 2,317,745 | 9,079,743 (35.2) | 168,599 (52.5) | 1,973 (52.4) | 11969 (45.8) |
|  | Extend 150 bp in each direction | 2,317,745 | 10,691,179 (41.5) | 184,175 (57.4) | 2,157 (57.3) | 13989 (53.5) |
|  | Extend 200 bp in each direction | 2,317,745 | 11,901,362 (46.2) | 196,938 (61.3) | 2,294 (60.9) | 15473 (59.2) |
|  | Extend 300 bp in each direction | 2,317,745 | 13,728,507 (53.3) | 217,011 (67.6) | 2,527 (67.1) | 17533 (67.1) |
| 450k array | - | - | 449,776 (1.7) | 36,780 (11.4) | 481 (12.7) | 770 (2.9) |
| EPIC array | - | - | 809,576 (3.1) | 65,417 (20.3) | 803 (21.3) | 1191 (4.6) |
| MeDIP-seq (hypermethylated markers) | - | - | 351,318 (1.3) | 30,716 (9.5) | 64 (1.7) | 1270 (4.9) |

\*Coverage of CpGs is defined as all CpGs (n = 25,746,133) annotated by hg38, not including chromosome X/Y/M and SNPs. Fragments of RRBS are filtered by the length $\geq 30$ bp; Cell type-specific markers (n = 320,838) are identified from 205 samples from a human methylation atlas; Coverage of the top 100 cell type-specific markers, the most differentiated 100 markers (n = 3,763) across 39 cell types from a human methylation atlas were used; 450k/EPIC array, not including chromosome X/Y/M, common SNPs, and ambiguously annotated CpGs.

An *in silico* analysis showed that the 100 bp regions flanking CCGG motifs cover 9 million CpGs with the highest yield closest to CCGG and 46% of CpGs in cell type-specific markers (Fig. 2a, Extended Data Fig. 1a-c). In contrast, other methylation profiling methods, including methylation microarrays, RRBS^13–16^, and methylated DNA immunoprecipitation sequencing (MeDIP-Seq)^18^ cover 3%-8% of those CpGs^11^. CCGG flanks also cover 36%-37% of CpGs from the machine learning (ML) classifiers based on methylation array data (central nervous system [CNS] or TCGA)^6,8,19^ (Fig. 2b, see Methods). CCGG flanks cover more enhancers, CpG shores/shelves, open sea, and introns than other methods (Extended Data Fig. 1d).

**Fig. 2:**
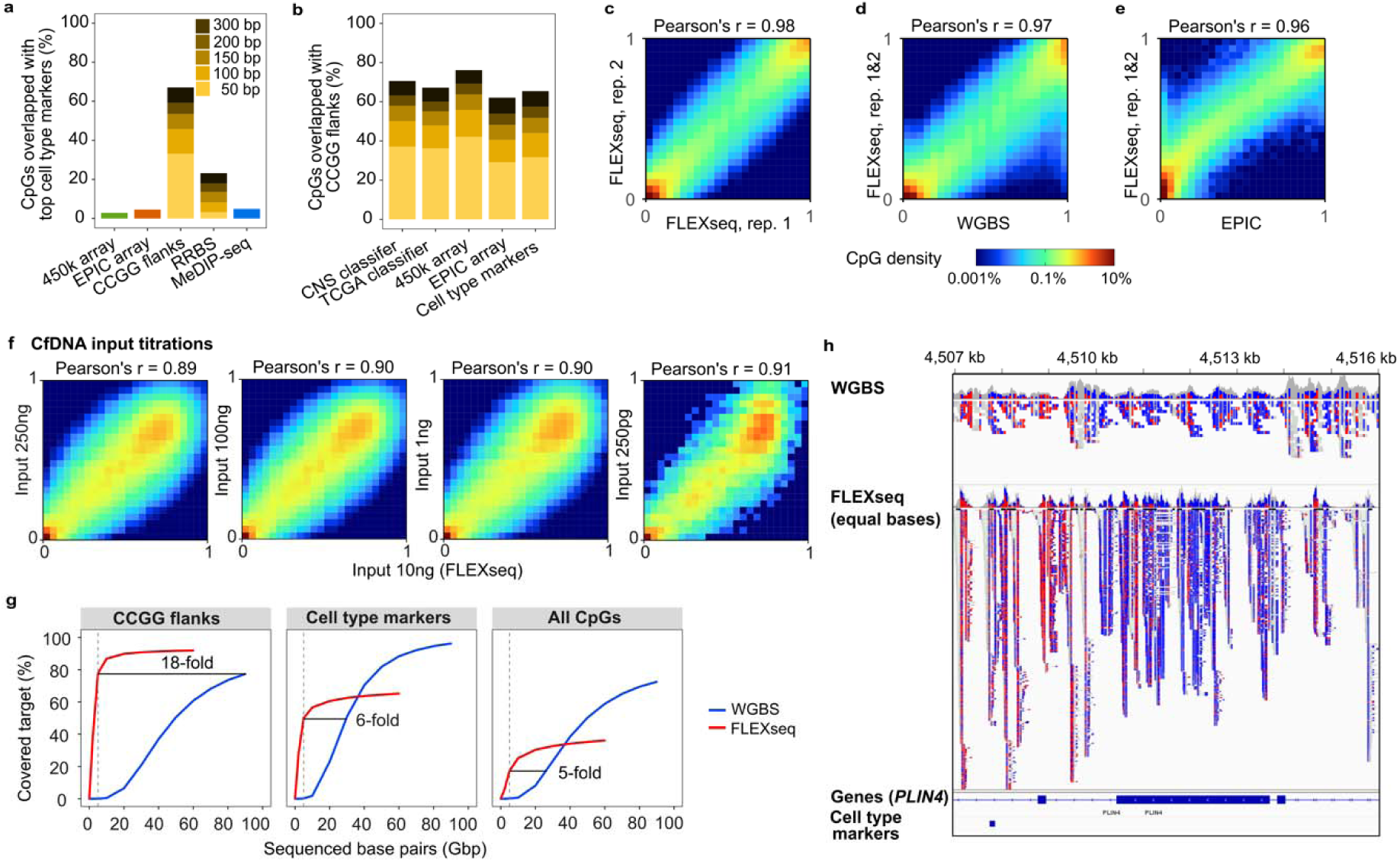
FLEXseq enrichment improves coverage at low methylation bias. **a,** Theoretical coverage of cell type markers^11^ (top 100 markers of each cell type) that overlap with different methods. **b,** Theoretical coverage of CpGs from CCGG flanks overlapped by different markers. The CNS classifier indicates the 32,000 most differentiated probes from the CNS tumor and control references^8^, and the TCGA classifier indicates the 60,000 most differentiated probes from TCGA tumor and control references^6,19^. **c,** Correlation of methylation beta values between inter-run replicates (Pearson’s r = 0.98). **d,** Correlation of beta values between FLEXseq and the WGBS gold standard (Pearson’s r = 0.97). **e,** Correlation between EPIC array and FLEXseq (Pearson’s r = 0.96). **f**, CfDNA input titrations sourced from a pleural fluid sample (BF3713) and assayed by FLEXseq. All non-10 ng inputs were correlated with 10 ng input (Pearson’s r ≥ 0.89). **g,** Enrichment of CCGG flanking 50 bp regions, the top 250 cell type-specific markers across different cell types^11^, and all CpGs with coverage 10x for WGBS (blue) and FLEXseq (red). Fold enrichment is based on 5Gbp of WGBS and FLEXseq. **h,** A representative area compares equal base pairs (30 Gbp) between WGBS and FLEXseq. Unmethylated (blue) and methylated (red) CpGs are highlighted. Gene and cell type marker tracks are shown at the bottom.

#### FLEXseq - design

We designed FLEXseq to target the CCGG flanks, output accurate methylation data, and be compatible with fragmented DNA input. FLEXseq involves two ligations and a double-stranded cut between the ligations, followed by methylation conversion (Fig. 1a). In the critical first step, both sides of input DNA molecules ligate on a semi-permissive adapter (Adapter A) that will serve as a blocker for untargeted background molecules and a primer landing site for targeted molecules. We then used the MspI nuclease to make double-stranded cuts, creating new and unblocked DNA ends. After an end repair, these newly exposed ends at the cut site are targeted through a second ligation with a second adapter (Adapter B) that is also needed for sequencing library formation. Because each of the two adapters at opposite ends has a necessary primer landing site (i.e. A+B are both needed), only targeted and cut DNA molecules are eventually sequenced while non-target DNA is ignored. Sequencing excludes non-target DNA with two A adapters or two B adapters. All DNA molecules have a free end associated with Adapter B that allows PCR duplicate removal and the potential of fragmentomics^20–23^. The distinct free end is a major advantage compared to most profiling methods besides WGBS. We then used enzymatic conversion^24^ instead of bisulfite conversion to avoid breaking the adapter-ligated molecules. Conversion is followed by amplification and sequencing of the target molecules.

#### FLEXseq - analytical performance

To assess reproducibility and bias in the methylation output, we correlated K562 replicates and compared them with WGBS as the gold standard. Two inter-run replicates were highly correlated (Pearson’s r = 0.98, Fig. 2c), demonstrating reproducibility. FLEXseq was highly correlated with WGBS, demonstrating biological fidelity (Pearson’s r = 0.97, Fig. 2d), and methylation array (Pearson’s r = 0.96, Fig. 2e). Three representative FFPE tissue samples were also correlated between FLEXseq and the EPIC version 2 (EPICv2) array (Pearson’s r = 0.93-0.95, Extended Data Fig. 2a-b). Moreover, FLEXseq had similar or higher correlations with WGBS than the public RRBS^25^ and XRBS^17^ data (Extended Data Fig. 2c). These high correlations suggest that FLEXseq data is compatible with past datasets such as ones derived from the microarrays.

We assessed the DNA input limit of detection by titrating cfDNA from pleural fluid (BF3713) and gDNA (K562). CpG methylation from titrated cfDNA was correlated to 10 ng input and had similar coverage down to 1 ng (Fig. 2f). Further dilutions down to 250 pg decreased CpG coverage, but bias in methylation remained low (Pearson’s r ≥ 0.90). Similarly, there was high correlation between 1-50 ng of FFPE tissue DNA (Pearson’s r ≥ 0.90, Extended Data Fig. 2d) and 1-100 ng K562 gDNA (Pearson’s r = 0.98, Extended Data Fig. 2e). In contrast, the EPICv2 methylation array titrations of K562 gDNA had a weaker correlation (Pearson’s r = 0.82) between 250 ng and 100 ng (Extended Data Fig. 2f).

The median on-target rate of K562 DNA input titrations was 96% (IQR 95%-96%) based on reads starting with (C/T)GG. Each on-target read is guaranteed to contain methylation data based on the first position. This high on-target rate yields an 18-, 6-, and 5-fold enrichment of CCGG flanks, cell type markers, and all CpGs compared with WGBS on a nucleotide basis (Fig. 2g). FLEXseq had the deepest coverage around the targeted CCGG cut sites (Fig. 2h, Extended Fig. 1c).

### 2. Analysis - copy number aberrations (CNA)

FLEXseq provided chromosomal copy number output after normalizing against control diploid references from CSF cfDNA. FLEXseq and whole genome sequencing (WGS) results were comparable, except in one case (Extended Data Fig. 3).

We detected tumor DNA by interpreting deviations from diploid as CNAs and indicators of clonal aneuploidy. We also estimated tumor purity based on the difference between the log2 ratios of chromosomal segments and the baseline at gains and losses, as previously described^26,27^.

### 3. Analysis - deconvolution

Studies have shown that tumor classification accuracy decrease with tumor purity <50-70% using machine learning for microarray data^28^, limiting the approach’s broader use in small tissue specimens and liquid biopsies. To overcome this limitation, we developed an orthogonal classifier based on the deconvolution of cell type proportions using cell type-specific markers.

#### Deconvolution of in silico titrations

To assess the performance of deconvolution in silico, we subsampled WGBS data^11^ from purified cell types and intersected them with CCGG flanks to create seven titrations (Table 2). We trialed two deconvolution approaches to predict cell type proportions for each titration: i) CelFiE CpG-level deconvolution^29^ using the top 30 cell type-specific markers that excluded the titration data, and ii) UXM fragment-level deconvolution using the top 250 markers. Both methods yielded similar predicted and actual proportions (root-mean-square error [RMSE] ≤ 0.01, Fig. 3a), with UXM fragment-level deconvolution remaining accurate down to 0.3% purity of target cell types.

**Fig. 3:**
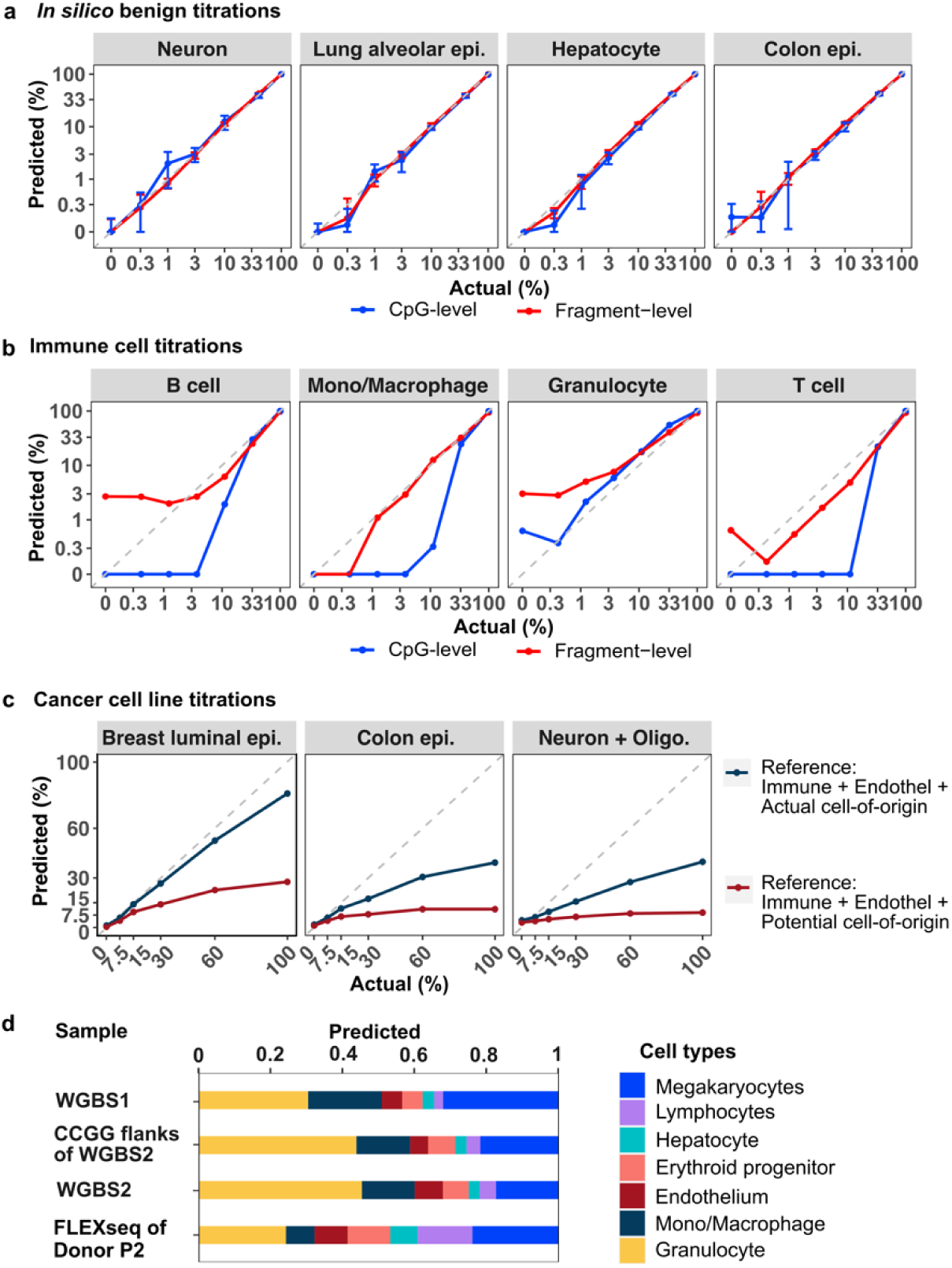
Deconvolution of *in silico* titrations, physical DNA titrations, and plasma. **a,** Cell type proportions deconvoluted from *in silico* mixtures using the WGBS references (WGBS reads intersected with CCGG flanks). CpG- (blue) and fragment-level deconvolution methods (red) are shown. Error bars indicate the SD at each titration level. **b,** Deconvolution of gDNA of B cells, monocytes, neutrophils, and T cells titrated into mixtures of three immune cell types, respectively. **c,** Fragment-level deconvolution of BRCA (tumor cell-of-origin, breast luminal epithelium), COAD (tumor cell-of-origin, colon epithelium), and GBM (approximate tumor cell-of-origin lineage, neuron and oligodendrocyte) cell line DNA titrated into mixtures of the same four immune cell types in (a). The dark blue line indicates deconvolution with references exclusive to the approximate tumor cell type and background immune cells. The dark red line indicates deconvolution with a broader set of reference cell types encompassing common brain metastases. **d**, Deconvolution of healthy plasma samples from Loyfer et al. (WGBS1), Gao et al. (WBGS2), WBGS2 intersected with CCGG flanks to simulate FLEXseq data, and FLEXseq from a healthy donor (P2).

**Table 2.** DNA mix-in titrations of four immune cell types and tumor cell lines in mixtures of those four immune cell types.

| Titration | B cell (%) | T cell (%) | Mono/Macrophage (monocyte, %) | Granulocyte (neutrophil, %) | Tumor cell (%) |
| --- | --- | --- | --- | --- | --- |
| **Immune cell DNA titration** |  |  |  |  |  |
| Bcell100 | 100 | 0 | 0 | 0 | 0 |
| Bcell33 | 33.33 | 22.22 | 22.22 | 22.22 | 0 |
| Bcell11 | 11.11 | 29.63 | 29.63 | 29.63 | 0 |
| Bcell3 | 3.7 | 32.1 | 32.1 | 32.1 | 0 |
| Bcell1 | 1.23 | 32.92 | 32.92 | 32.92 | 0 |
| Bcell0.3 | 0.41 | 33.2 | 33.2 | 33.2 | 0 |
| Bcell0 | 0 | 33.33 | 33.33 | 33.33 | 0 |
| Monocyte100 | 0 | 0 | 100 | 0 | 0 |
| Monocyte33 | 22.22 | 22.22 | 33.33 | 22.22 | 0 |
| Monocyte11 | 29.63 | 29.63 | 11.11 | 29.63 | 0 |
| Monocyte3 | 32.1 | 32.1 | 3.7 | 32.1 | 0 |
| Monocyte1 | 32.92 | 32.92 | 1.23 | 32.92 | 0 |
| Monocyte0.3 | 33.2 | 33.2 | 0.41 | 33.2 | 0 |
| Monocyte0 | 33.33 | 33.33 | 0 | 33.33 | 0 |
| Neutrophil100 | 0 | 0 | 0 | 100 | 0 |
| Neutrophil33 | 22.22 | 22.22 | 22.22 | 33.33 | 0 |
| Neutrophil11 | 29.63 | 29.63 | 29.63 | 11.11 | 0 |
| Neutrophil3 | 32.1 | 32.1 | 32.1 | 3.7 | 0 |
| Neutrophil1 | 32.92 | 32.92 | 32.92 | 1.23 | 0 |
| Neutrophil0.3 | 33.2 | 33.2 | 33.2 | 0.41 | 0 |
| Neutrophil0 | 33.33 | 33.33 | 33.33 | 0 | 0 |
| Tcell100 | 0 | 100 | 0 | 0 | 0 |
| Tcell33 | 22.22 | 33.33 | 22.22 | 22.22 | 0 |
| Tcell11 | 29.63 | 11.11 | 29.63 | 29.63 | 0 |
| Tcell3 | 32.1 | 3.7 | 32.1 | 32.1 | 0 |
| Tcell1 | 32.92 | 1.23 | 32.92 | 32.92 | 0 |
| Tcell0.3 | 33.2 | 0.41 | 33.2 | 33.2 | 0 |
| Tcell0 | 33.33 | 0 | 33.33 | 33.33 | 0 |
| **Cancer cell line DNA titration** |  |  |  |  |  |
| BRCA100 | 0 | 0 | 0 | 0 | 100 |
| BRCA60 | 6 | 14 | 6 | 14 | 60 |
| BRCA30 | 10.5 | 24.5 | 10.5 | 24.5 | 30 |
| BRCA15 | 12.75 | 29.75 | 12.75 | 29.75 | 15 |
| BRCA7.5 | 13.875 | 32.375 | 13.875 | 32.375 | 7.5 |
| GBM100 | 0 | 0 | 0 | 0 | 100 |
| GBM60 | 10 | 10 | 10 | 10 | 60 |
| GBM30 | 17.5 | 17.5 | 17.5 | 17.5 | 30 |
| GBM15 | 21.25 | 21.25 | 21.25 | 21.25 | 15 |
| GBM7.5 | 23.125 | 23.125 | 23.125 | 23.125 | 7.5 |
| COAD100 | 0 | 0 | 0 | 0 | 100 |
| COAD60 | 10 | 10 | 10 | 10 | 60 |
| COAD30 | 17.5 | 17.5 | 17.5 | 17.5 | 30 |
| COAD15 | 21.25 | 21.25 | 21.25 | 21.25 | 15 |
| COAD7.5 | 23.125 | 23.125 | 23.125 | 23.125 | 7.5 |

#### Deconvolution of DNA titrations and plasma

We continued with physical titrations by mixing DNA from purified primary immune cell types (Table 2). Fragment-level deconvolution detected target cell types down to 1% purity, and we used this approach for all further deconvolutions (Fig. 3b). We then deconvoluted the titrations of DNA from cancer cell lines into DNA from immune cell mixtures. We found that the deconvolution underestimated the tumor cell type proportion, but the rank order of the proportions was reliable (Fig. 3c). We then deconvoluted plasma cfDNA from healthy donors^30^ using WGBS and FLEXseq data (Fig. 3d). Cell type proportions were similar between WGBS and WGBS intersected with CCGG flanks (RMSE = 0.02).

## Discussion

This paper introduces FLEXseq, a methylation enrichment profiling assay, and demonstrates its clinical potential through copy number detection across different liquid biopsies and FFPE tissues. FLEXseq achieves an 18-fold enrichment covering CCGG flanking regions with an on-target rate of 96%-98%, guaranteeing methylation data from nearly every sequenced DNA molecule. FLEXseq is highly concordant with the WGBS gold standard, accurately reflecting biological truth. Its genome-wide coverage yields low-noise copy number plots. Its broad coverage allows for integration with public datasets (WGBS and microarray references) for tumor classification.

We used TET, BGT, and APOBEC enzymatic conversions^24^, but alternatives include other non-destructive conversion strategies such as TAPS^31^, DM-seq^32^, or SEM-seq^33^. Bisulfite conversion is possible, albeit with DNA loss. Because the enrichment occurs before PCR, we expect FLEXseq to be compatible with direct methylation readout using nanopore sequencing^34^. Moreover, the design of FLEXseq is not limited to the MspI nuclease, which is restricted to CCGG motifs. Other nucleases like CRISPR-Cas9 can target alternative flanking regions.

There are numerous methylation profiling assays, but each has limitations in clinical use. <u>WGBS</u> is the gold standard, but its high sequencing and computational costs bar large-scale studies or clinical testing. <u>Methylation microarrays</u> are commonly used but require 250 ng of DNA input and cover only 2%-4% of all CpGs and 3%-8% of cell type markers (Fig. 2a), limiting deconvolution accuracy^11^. Building array reference sets is costly due to the need to scale to thousands of samples^6,8,19^. The EPICv2 array costs $265 per sample and $70 for FFPE repair. In this study, 160M reads amount to $114 per sample (NovaseqX 25B kit). As research tools, microarrays are restricted to human and mouse genomes. <u>RRBS</u>^13–16^ is effective for CpG islands but covers fewer enhancers and cell type-specific markers (Extended Data Fig. 1). <u>XRBS</u>^17^ targets CCGG flanks like FLEXseq but was not designed for fragmented DNA found in clinical specimens. <u>MeDIP-seq</u> targets methylated cytosines but imprecisely detects hypomethylated markers, encompassing 98% of cell type-specific markers.

As a cost-efficient yet broad profiling technology, FLEXseq opens future possibilities. By preserving one free end on every DNA molecule, it has the potential to be utilized in fragmentomics^20–23^. FLEXseq captures half of the known methylation aging markers associated with epigenetic clocks^9,35^ and is poised to broaden the identification of aging markers. FLEXseq can be used to explore the methylome in non-human organisms^35^. We expect to phase genotypes and mutations with the methylation status in over 95% of reads. Finally, our data is compatible with metagenomics, which could potentially provide information relevant to infectious diseases^36^.

FLEXseq has limitations like any method. Being a broad profiling assay, it does not achieve the same depth as targeted panels using the same amount of sequencing. Target panels also require prior knowledge of the markers, a major limitation when only 40 of the >400 human cell types have purified references^11,37^. Our deconvolution process is currently limited to 40 pure cell type references, whereas RNA cell atlases show that >400 cell types exist^37^. However, purified cell type references will only increase over time, especially with the advent of single-cell sequencing^38^ and the usage of pseudo-bulk data.

## Data availability

Raw sequencing data is available for all cell lines and samples from healthy donors who gave informed consent for genomic data sharing. FASTQ files have been deposited with SRA under Bioproject PRJNA1125505. DNA methylation data across all samples is available in deidentified PAT files to ensure fragment-level read information is preserved while protecting the privacy and confidentiality of patients. PAT files, methylation calls data, copy number plots, and *t*-SNE plots are available under Zenodo https://doi.org/10.5281/zenodo.12668041. The data for figures are available at GitHub https://github.com/GuLabDNA/MethylSeq.

## Code availability

The R codes for figures are available at GitHub https://github.com/GuLabDNA/MethylSeq.

## Acknowledgments

This work was funded by an NIH K08 (CA230156) grant, a Burroughs-Wellcome CAMS Award to WG, a DoD grant to BH, and an NIH U54 Brain MetNet fellowship grant to JY. We thank the members of the Stanford Clinical Molecular Genetic Pathology, Cytology, and Flow Cytometry Laboratories and UCSF Clinical Immunology and Hematology Laboratories for helping to save residual specimens. We thank members of Stanford Molecular Pathology and Clinical Genomics, particularly Chandler Ho, Linda Liao, Tina Tan-Pruitt, Diwash Jangam, Eula Fung, James Zehnder, and Jean Oak, for help with specimen workflow, automation, sequencing, and informatics. We thank Tong Wang for critical feedback. We thank members of the Howitt, Lowe, and Hayden-Gephart laboratories for their support. We thank The University of Chicago Genomics Facility (RRID: SCR_019196) for methylation array data and Novaseq sequencing and the Pathology Molecular Core for sequencing.

## Contributions

J.Y., L.S.A., Y.Y.Y., and W.G. conceptualized and designed the study. L.S.A., Y.Y.Y., and W.G. prepared and sequenced the samples. J.Y. and W.G. wrote the bioinformatic pre-processing pipeline, L.S.A. performed the quality control analysis, and J.Y. performed the bioinformatics analysis and visualization. L.S.A. and W.G. collected clinical information from medical charts. J.Y., L.S.A., and W.G. wrote the manuscript. All authors revised the manuscript and approved it for publication.

## Extended Data Figures

**Extended Data Fig. 1:**
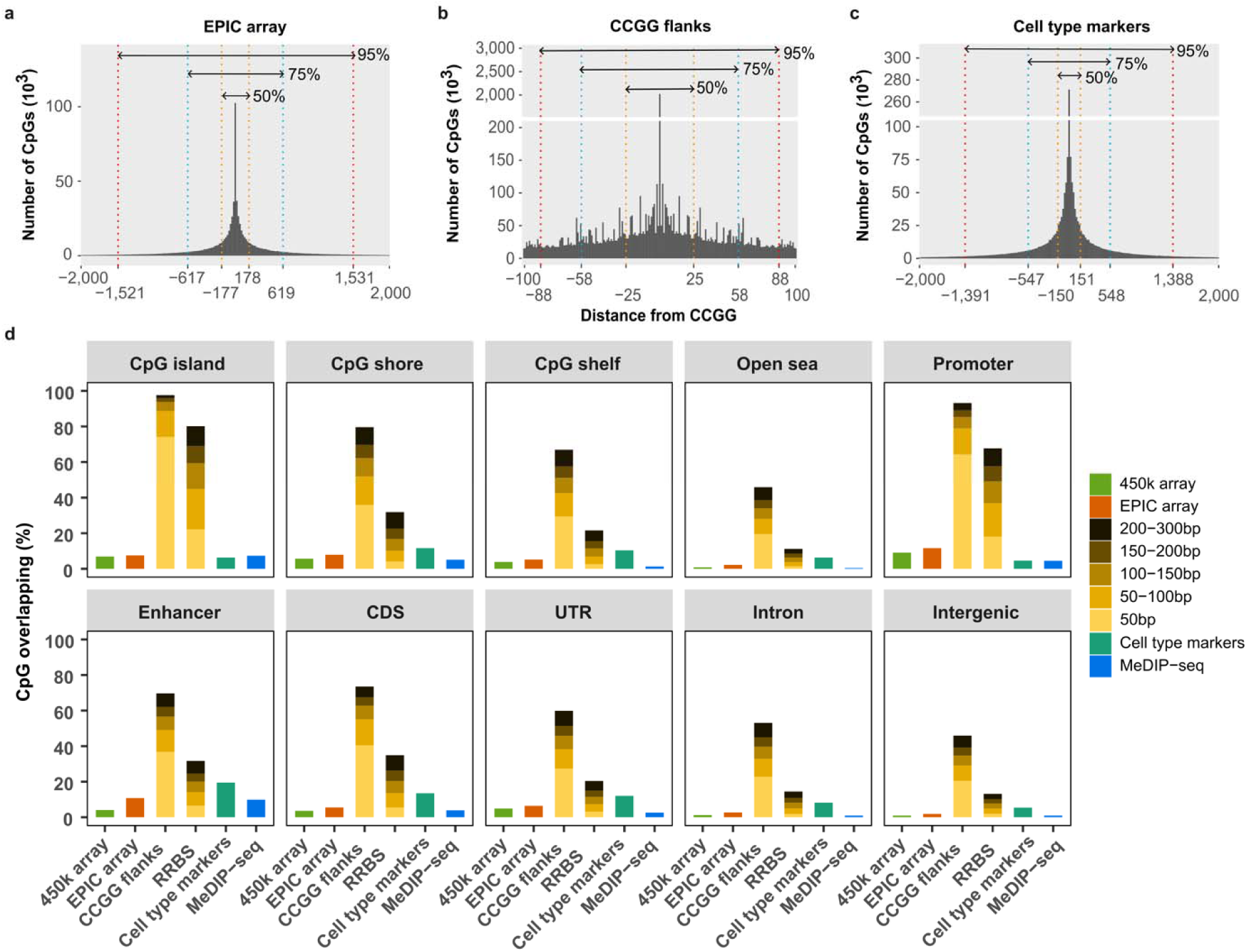
*in silico* coverage of CpGs across genomic regions by different methods **a**, CpG percentages across genomic regions by different methylation detection methods. CCGG flanks are the 50-300 bp regions flanking CCGG motifs. RRBS regions are two CCGG motifs within a maximum distance of 50-300 bp. Cell type markers are defined across 39 purified cell types from a human DNA methylation atlas. MeDIP-seq markers are defined as CpGs in hypermethylated regions derived from the above cell type-specific markers. The ranges between the orange, blue, and red dashed lines indicate the 50%, 75%, and 95% distribution of CpGs, respectively. CDS, coding sequence; UTR, untranslated region, including 5’ and 3’ UTR. **b,c,d**, Theoretical distance from the CCGG motifs for all CpGs covered by EPIC array, CCGG flanks, and cell type markers.

**Extended Data Fig. 2:**
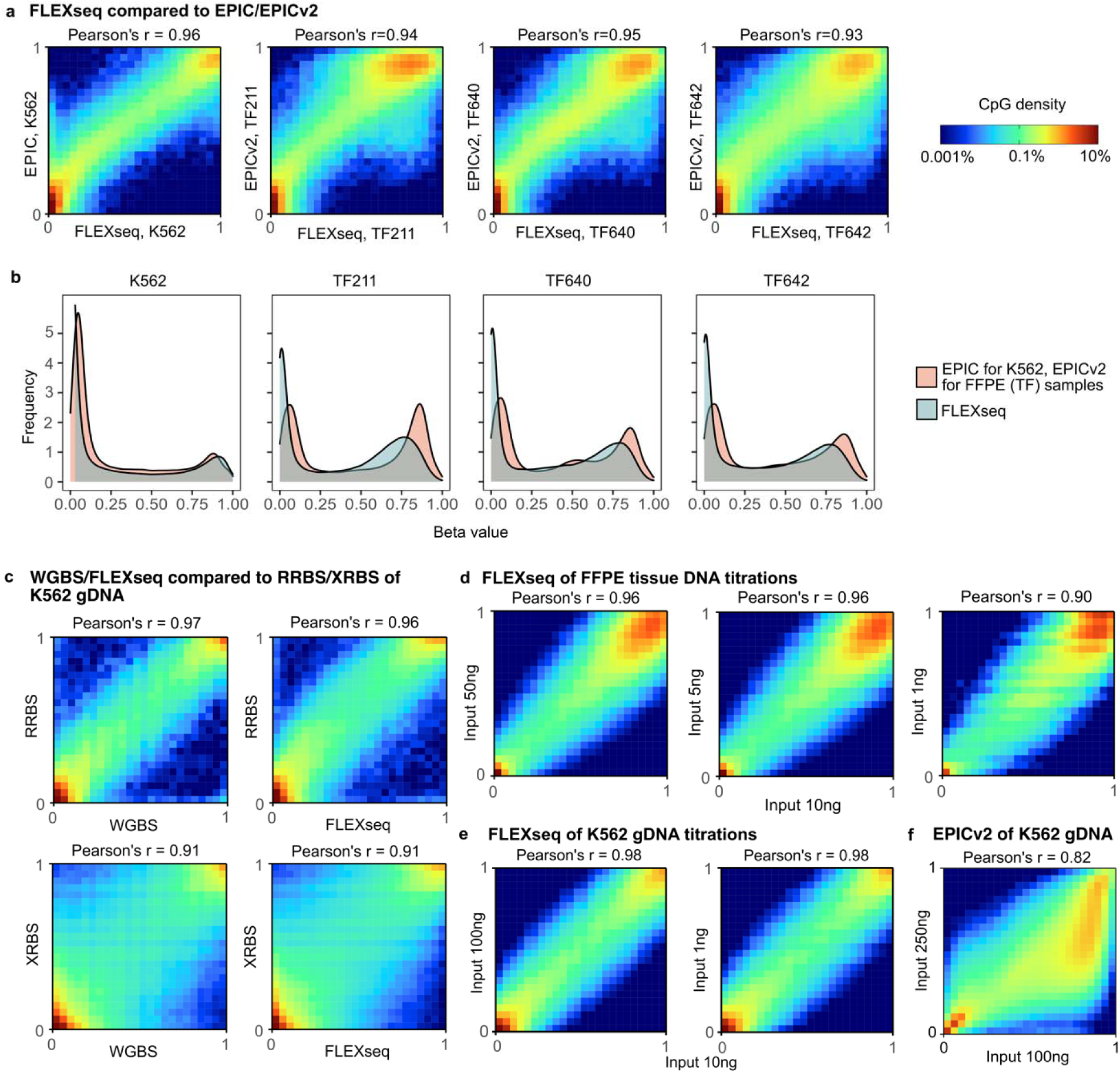
Correlations of methylation beta values across different methods, sample types, and DNA inputs. **a,** FLEXseq is highly correlated with the EPIC array, as observed in K562 and three FFPE tumor tissue samples (TF211, TF640, and TF642). **b,** Density plots of beta values of the same four samples in (a). **c,** Correlation between WGBS, FLEXseq, and RRBS/XRBS of K562 gDNA. All correlations are based on individual CpG methylation beta values. **d,** FFPE tissue (TF640) DNA input titrations assayed by FLEXseq. Inputs of 1, 5, and 50 ng were correlated with 10 ng input (Pearson’s r >= 0.90), though the number of covered CpGs decreased from 5 million with 10 ng to 0.2 million with 1 ng. **e**, K562 gDNA input titrations assayed by FLEXseq. Inputs of 1 ng and 100 ng were correlated with 10 ng input (Pearson’s r = 0.98). **f**, K562 gDNA input of EPICv2 array at 100 ng was less correlated with the recommended input of 250 ng (Pearson’s r = 0.82).

**Extended Data Fig. 3:**
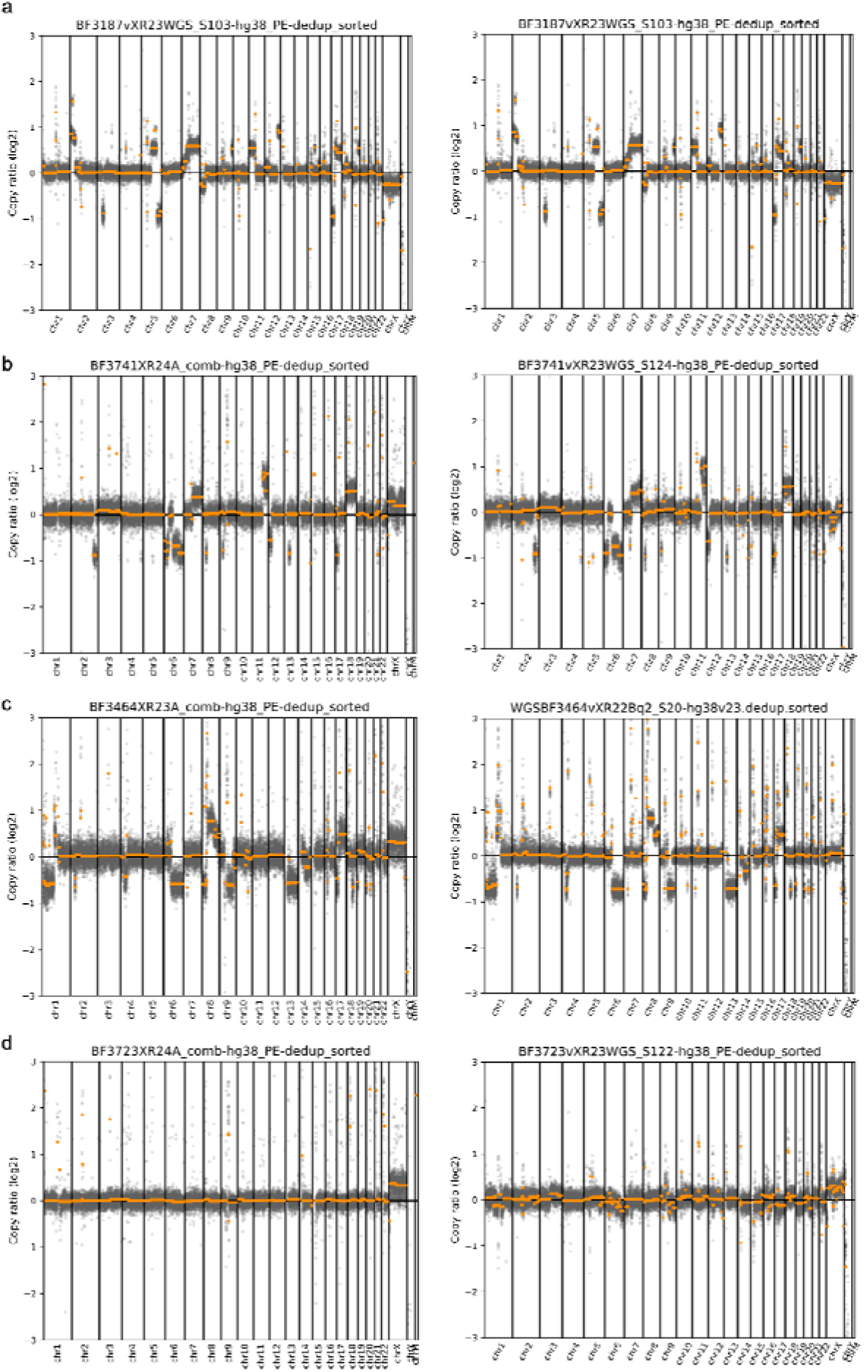
Copy number analyses from FLEXseq and WGS. Paired copy ratio plots between FLEXseq (left) and WGS (right) are shown. Chromosomal regions had comparable results, excluding the sex chromosomes (chrX and chrY), which were not adjusted by sex. Two are tumor positives from CSF cfDNA, BF3187 (**a**) and BF3741 (**b**), and one, BF3464 (**c**), was a positive from saline wash fluid cfDNA of a liver fine-needle aspiration (FNA) biopsy. **d**, BF3723 was the only case overall with CNA-positive in WGS data but not from FLEXseq. The full set of plots is available (see Data Availability).

## Methods

### Ethics statement

Body fluid and FFPE tissue samples from Stanford Healthcare were originally processed by the Stanford Pathology and Clinical Laboratories as part of routine clinical testing, and residual material was retrospectively enrolled through a waiver of consent in a protocol approved by the Stanford Institutional Review Board (IRB 58461). Plasma P2 control was collected under informed consent in a protocol approved by the Stanford Institutional Review Board (IRB 71230). IRB approvals and sample collection details from the UCSF Medical Center are detailed in past publications^26,27,36^.

### Sample collection

We used CSF and non-CSF body fluid samples derived from previous studies^26,27,36^ and new collections. New body fluid samples were consecutively from Cytology and Flow Cytometry Laboratories at Stanford Healthcare between 2020 and 2023. Specimens from cytology were spun following the clinical standard operating procedure (∼800-1600g for 10 minutes or by gravitational settling). The supernatant was then pipetted or decanted and then stored at refrigeration until it was processed for cfDNA. Body fluids specimens from flow cytometry were kept at 4°C for up to 4 weeks and then processed. For fine needle aspirates (FNA), the clinical protocol used normal saline fluid from needle rinses or specimens placed directly into normal saline. We also selected 3 cases with FFPE tissue DNA from patients with a clear pathological diagnosis. The tissue was across various body sites to check the feasibility of FLEXseq.

The clinical and demographic information of all patients was reviewed retrospectively from medical records. Pathological diagnosis serves as the gold standard for tumor classification.

### In silico pre-design analysis

#### *In silico* CCGG flanks, cell type, array, RRBS, and MeDIP-seq markers

For the *in silico* analysis before designing FLEXseq, we intersected CpGs from cell type markers (Fig. 2a) and CCGG flanks (Fig. 2b) with other markers and methods. Cell type markers are the top 100 markers for all 39 cell types; see the section ‘*Genomic segmentation and identification of cell type-specific markers*’ below for details. For CpGs on the 450k and EPIC arrays, we downloaded the annotation files from https://zwdzwd.github.io/InfiniumAnnotation. For CCGG flanks, we produced all CCGG motif coordinates in a BED format and expanded the CCGG motifs to specific lengths with ‘bedtools slop -b 50 (100/150/200/300)’. RRBS regions are defined as two CCGG motifs within a maximum distance of 50-300 bp of each other, with the command ‘bedtools merge -d 50 (100/150/200/300)’. The RRBS regions shorter than 30 bp were dropped. For the MeDIP-seq markers, we used only cell type-specific hypermethylated markers since the assay pulls down on methylated markers. For the CNS and TCGA classifier markers, we used the top 32,000 and 60,000 probes, respectively.

#### Annotation of promoters, enhancers, and other methylation regions

We divided the CpG regions according to genomic localization: CpG islands; shores, regions up to 2 kb from CpG island; shelves, regions from 2 to 4 kb from CpG island; and open sea, the rest of the genome. Promoters were defined as 1 kb upstream and downstream of all transcription start sites (downloaded from UCSC) of protein-coding genes. Enhancers were downloaded from Fantom5. Introns, coding exons, 5’ untranslated regions (UTR), and 3’ UTRs were parsed from RefSeq annotation. Intergenic regions were the regions excluding promoters, enhancers, exons, untranslated regions, and introns within protein-coding genes. Each called CpG site is counted once: the overlap of a genomic region (promoters, enhancers, coding sequences, untranslated regions, and introns) excludes all previously overlapped sites starting from the promoter. The denominator is the total CpGs across genomic regions, and the numerator is the CpGs covered by methylation detection methods.

### Sample DNA extraction

#### Cell lines

DNA from MCF-7 (ACC 115, BRCA), CL-40 (ACC 535, COAD), and DBTRG-05MG (ACC 359, GBM) were obtained from DSMZ (Braunschweig, Germany), and DNA from K562 (E4931) was obtained from Promega.

#### Primary cells

Purified immune cells (purity above 95% by flow cytometry) were obtained from IQ Biosciences, and dissociated cells were extracted using the Quick-DNA Miniprep kit (D3020, Zymo Research).

#### Fresh body fluid

Up to 1 mL of DNA was extracted by the Maxwell RSC ccfDNA Plasma Kit (AS1480, Promega) with approximately 47 µL output. For inputs greater than 1 mL, we used the RSC ccfDNA LV kit instead (AS1480, Promega). Extracted DNA was quantified with Qubit, excluding CSF cfDNA samples with a white blood cell count < 1000 cells/µL.

#### FFPE

FFPE blocks or previously extracted and frozen (-80°C) FFPE tissue DNA were used. DNA extraction from FFPE blocks was performed using the Maxwell RSC FFPE Plus DNA kit (AS1720, Promega) or QIAamp DNA FFPE Tissue Kit (PN 56404, Qiagen). The clinical molecular pathology lab found no difference in output between the two kits. Extracted DNA was quantified on a spectrometer (Nanodrop, Thermo Fisher) or Qubit (Thermo Fisher). Frozen DNA was stored in DNA Lobind tubes at -20°C.

### Sequencing library preparation

#### Shearing

DNA from cell lines, immune cells, and FFPE DNA were sheared with acoustics using a 96 AFA-TUBE TPX plate (PN 520291, Covaris) set at 400-500 bp.

#### Sequencing

Sequencing was performed on an Illumina Novaseq 6000 using 100 cycle S2 kits configured as paired-end 2x50 bp. To address the decrease in nucleotide diversity due to the C to T conversions, we spiked in 10-15% PhiX sequencing library DNA (Illumina) or whole genome libraries. Samples were only included in library pooling if their qPCR curve was ≤14 Ct (estimated to the nearest 0.5). Library pooling involves equivolume pooling because the PCR amplification is to saturation, and concentrations are determined by the equimolar primer input.

### Sequencing alignment and preprocessing

Paired-end reads were quality and length trimmed with cutadapt v.3.5 with the following parameters: -a NNAGATCGGAAGAGC -A NNAGATCGGAAGAGC --minimum-length 25. High-quality sequencing reads were then aligned to the hg38 reference genome using Bismark v.0.23.0. Sorted alignments were further processed to only maintain uniquely mapped read pairs with alignment score ≥-5 (--score_min L,0,-0.2), ignoring the quality values of individual bases during the alignment process, and discarding reads with multiple alignments.

Incomplete enzymatic conversion affected large portions of the same read, allowing us to filter out reads with unmethylated cytosine in the non-CpG context with the filter_non_conversion function from Bismark. Using this filter, we found that FFPE samples have a higher filter rate (5.1%) than non-FFPE samples (0.9% for gDNA and 0.5% for cfDNA). The detection of non-converted DNA molecules highlights an advantage of base-resolution sequencing to correct for incomplete conversions.

After filtering non-CG converted and duplicated reads, we used the bismark_methylation_extractor function to extract the methylation calls with the option -- bedGraph. The methylation calls of a CpG dinucleotide entity from both top and bottom strands, staggered by 1 bp, were merged using the coverage2cytosine function with the parameter --discordance 50, to increase coverage per paired CpG location and reduce the memory burden in downstream analyses. Next, all single-nucleotide polymorphism (SNP) positions (dbSNP153Common) were removed for deidentification using bedtools.

We also used the bam2pat function from wgbs_tools^39^ to convert bam files into PAT and BETA files for deconvolution, keeping reads covering at least three CpG sites (-- min_cpg 3). The PAT files preserved fragment-level data and were deidentified by removing SNPs using the mask_pat function. All deidentified PAT files from each sample are available and used for downstream analyses (see Data Availability).

### Obtaining and processing external data from alternative methods

Outside data was obtained and pre-processed as follows: To correlate FLEXseq with WGBS, we downloaded the WGBS data for K562 cells (SRR4235743) as raw FASTQ files. The paired-end WGBS data was trimmed by 15 bp from both the 5’ end of Read 1 and the 3’ end of Read 2 (cutadapt -a AGATCGGAAGAGC -A AGATCGGAAGAGC -u - 15 -U 15). Other pre-processing steps were similar to FLEXseq. We also downloaded the RRBS data (GSM683856 and GSM683780) as raw FASTQ files. The single-end RRBS data was trimmed using trim_galore function from TrimGalore v.0.6.10, with parameters ‘--rrbs --phred64 --quality 33 --illumina’. Other pre-processing steps were similar to FLEXseq with single-end data. XRBS data (GSM4518657 and GSM4518657) derived from 10 ng DNA were downloaded as methylation metadata.

Our internal array data DNA from the FFPE tissue samples were processed by the University of Chicago Genomics Facility without DNA repair. External K562 methylation EPIC array data (ENCFF848ACR and ENCFF849HRI from ENCODE) was downloaded as IDAT files.

WGBS data of plasma cfDNA from 39 healthy individuals was obtained from Gao and colleagues^30^ (CRA001142). The raw FASTQ files were processed following a similar pipeline as FLEXseq but with a broader trimming with parameters ‘-u -10 -U 10’. We used the bismark_methylation_extractor function to get methylation calls for individual CpGs. Covered CpG sites were further intersected with CCGG flanks to align with FLEXseq data. To increase the sequencing coverage, we randomly merged 13 patients to create three combined samples with a median coverage of 10x.

### Analytical benchmarks

Benchmarks were primarily performed on sheared K562 DNA. The correlation of methylation profiles between different approaches was tested using overlapping CpG sites with ≥ 15 coverage. Pearson’s correlation coefficient was calculated using the cor.test R function with single-CpG resolution.

For the reproducibility analysis, we sequenced two K562 replicates at 20 ng gDNA input across different library preparation and sequencing runs.

For the enrichment analysis (Fig. 2g-h), we randomly subsampled the aggregated reads of K562 from WGBS (358,053,131 paired-end reads with a length of 2x125 bp) and FLEXseq (657,614,778 paired-end reads with a length of 2x50 bp) down to 20-900 million paired-end reads. To calculate the fold-enrichment, we excluded CpGs with coverage <10x from WGBS and FLEXseq, and then compared the target enrichment between them based on 5 Gbp data (50 million reads of 2x50 bp of FLEXseq and 20 million reads of 2x125 bp of WGBS). CpG sites intersected with the CCGG flanks and cell type markers (the top 250 UXM markers from a human DNA methylation atlas^11^,) were counted for comparisons.

To evaluate the performance of different DNA amounts, we titrated DNA inputs from 250 ng to 100, 10, 1, and 0.25 ng of one pleural fluid sample (BF3713, Fig. 2f) and tested their correlations based on overlapping CpGs sites with coverage ≥15x.

### Copy number analysis

We used CNVkit (v0.9.10)^40^ to analyze and visualize genome-wide copy numbers. Our inputs into CNVkit were Bismark/Bowtie 2 aligned BAM files that were deduplicated by Bismark based on end positions and fragment lengths (see above).

Proper normalization is critical to denoise copy number plots. We employed a cfDNA pooled reference to normalize FLEXseq regions for CNA calling. First, a healthy control plasma (P2) was used as an initial reference. We ran CNVkit on ‘wgs’ mode and used a hg38 flat file to split the genome into 50 kbp blocks. This reference was then used to generate CNR files of a randomized selection of four CSF cfDNA samples (BF3040, BF3222, BF3241, and BF3361) with no malignancies and >50 million reads. Noise in each sample was evaluated using the CNVkit metrics command, and one sample with high-level noise (BF3014, with a larger biweight midvariance, median absolute deviation, and IQR) was removed. The final pooled reference utilizing those four BAM files from CSF cfDNA was generated following the same steps as with the P2 reference. We did not track and normalize the sex of specimens, and therefore, copy numbers from the sex chromosomes are not reliable and subsequently ignored during analyses.

We then generated log2copy ratio plots for all body fluid and FFPE samples based on the pooled reference and visualized them across all bins. The gray points in the plots are binned ratios from CNR files and the orange lines are inferred discrete copy number segments derived from CNS files. We derived the segments based on the default Circular Binary Segmentation algorithm from the DNAcopy R package. Samples were considered CNA-positive if the copy ratio plot showed one or more significant CNAs, excluding sex chromosomes.

Tumor purity was estimated using the log2 copy ratio from segments larger than 10 Mbp by measuring the maximal deviation from the baseline, which was assumed to be diploid. Certain deletions or gains were assumed to be single copy changes (e.g., monosomy or trisomy). The following equation was used to determine the tumor purity as previously described^26,40^.

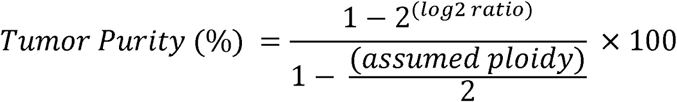

### In silico titrations

We mixed methylation profiles from the human DNA methylation atlas^11^ (WGBS) samples with different cell type compositions, including B cell, T cell, monocyte, granulocyte, and/or hepatocyte, colon epithelium, lung alveolar epithelium, neuron, oligodendrocyte, and endothelium. We simulated seven mixtures at proportions of 0%, 0.3%, 1%, 3%, 10%, 40%, and 100% of target cell types, conducting three mixtures with two replicates for each level. Merging, splitting, and mixing of reads were performed using wgbs_tools functions ‘merge’ and ‘mix_pat’. We then dropped the CpGs that are covered by WGBS but not CCGG motifs flanked by 50 bp to simulate the FLEXseq data. Matching references from the atlas were used, from which the individual mixed-in samples were excluded.

### In vitro DNA titrations

We extracted, sheared, and quantified DNA from four human immune cell types (B cell, T cell, monocyte, and neutrophil) as described above. DNA was diluted to 5 ng/μL each and then mixed at different proportions to simulate benign samples with a high immune background. Specifically, two background mixtures were made: i) 25% of each cell type and ii) 15% each of B cells and monocytes and 35% each of T cells and neutrophils. They were then titrated separately with DNA from three tumor cell lines: BRCA, COAD, and GBM (see details in the ‘*Sample DNA extraction’* section). Tumor cell line DNA was titrated from 100% down to final purities of 60%, 30%, 15%, and 7.5% (Table 2).

### Genomic segmentation and identification of cell type-specific markers

#### Identification of haplotypes

We used the wgbs_tools function ‘segment’ with 205 samples as references (grouped into 39 groups according to their methylation patterns) to segment the genome into 2,804,836 haplotypes that cover at least 3 CpGs (with parameters --min_cpg 3 -- max_bp 5000), excluding chrX, chrY, and chrM. Each haplotype should have homogeneous methylation levels across multiple consecutive CpGs and cell types. The average methylation per haplotype in a sample was calculated by summarizing methylated and unmethylated calls across the entire region and then calculating a single beta value (methylated counts divided by total counts). We further kept the haplotypes with a length of 10-2,000 bp, and CpGs in these haplotypes were intersected with the methylation array probes for computational inference.

#### Identification of cell type-specific markers

We performed a one-vs-all comparison to identify differentially methylated segments with a length of 10-2,000 bp for each cell type, using the wgbs_tools function ‘find_markers’ (parameters ‘--min_cpg 4 --min_cov 10 --delta_means .3 --pval .05’). From the initial list of segments representing cell type-specific markers, we then selected markers with the following criteria: i) the average methylation ≥0.66 of one cell type and <0.33 in all others, or vice versa, and ii) coverage of 30x or more (across all reads that cover CpGs of the marker). We selected the top 30 and 100 markers with the highest delta of beta values for each cell type (for cell types with more than 1,000 markers, we further used parameters ‘delta_means >0.4 and delta_quants >0.1’). Hypomethylated markers were defined based on the difference between the 75th percentile among the segment average methylation within the target samples and the 2.5th percentile among the background samples. Hypermethylated markers were defined based on the difference between the 97.5th percentile of the background and the 25th percentile within the target samples. All filtered and the top 100 markers were used for *in silico* analyses.

For cell type-specific markers across immune cells (B cell, T cell, monocyte, granulocyte) and/or breast luminal epithelium, colon epithelium, lung alveolar epithelium, neuron, oligodendrocyte, and endothelium, we divided the genome into segments with lengths of 10-2,000 bp, and then identified markers with parameters ‘--min_cpg 5 --min_cov 10 --delta_means .3 --pval .05’. The top 30 markers were used for further analyses. CpGs from those WGBS cell type references were filtered by intersection with CCGG flanks to align with FLEXseq’s coverage before finding markers.

### Deconvolution of cell types

We conducted deconvolution analyses using CpG-level CelFiE (expectation-maximization algorithm)^29^ and fragment-level UXM (non-negative least-squares algorithm)^11,39^. For the CpG-level deconvolution, the top 30 markers across different cell type references were identified and filtered with the criteria mentioned in the section ‘*Genomic segmentation and identification of cell type-specific markers*’ above. Methylated counts and total counts for each individual CpG within the marker were summed up to calculate the marker’s beta value.

For the fragment-level deconvolution, the top 250 unmethylated markers across different cell type references were used. Those markers were extracted directly from Loyfer et al^11^. We removed markers on the sex chromosomes and those that overlapped with common SNP positions. Different cell type references were used across various scenarios:

When deconvoluting *in silico* samples, we used 180 markers for CpG-level deconvolution and 1,500/1,750 markers (FLEXseq data covers ∼60% of them) for the fragment-level deconvolution. These markers not only came from four immune cell types (B cell, T cell, mono/macrophage, and granulocyte) and endothelium, but also from the specific tissue cells. We included hepatocyte for liver mixtures, colon epithelium for colon mixtures, lung alveolar epithelium for lung mixtures, and neuron and oligodendrocyte for brain mixtures. When deconvoluting physical immune cell titrations, we used 120 (CpG-level) or 1,000 (fragment-level) markers across the four immune cell types.

When deconvoluting tumor cell line DNA titrations, we used the fragment-level deconvolution with two reference sets: i) B cell (immune/B cell lymphomas), T cell (immune/T cell lymphomas), mono/macrophage (immune), granulocyte (immune), endothelial and smooth muscle cell (vasculature), and the specific tumor cell-of-origin (luminal and basal breast epithelium for breast carcinoma, colon and small intestine epithelium for colon carcinoma, or neuron and oligodendrocyte for primary CNS tumors and neuronal cells); or ii) additional nine cell types associated with common epithelial malignancies, including lung alveolar and bronchial epithelium (lung carcinoma), hepatocyte (hepatocellular carcinoma), pancreatic ductal epithelium (pancreatic carcinoma), ovarian and endometrium epithelium (ovarian and uterine carcinomas), gastric epithelium (gastric carcinoma), kidney epithelium (renal cell carcinoma), and bladder epithelium (bladder carcinoma).

### Statistical analysis

Wilcoxon rank-sum test was used to test the distribution differences of continuous variables between groups. Adjusted *P* value was used for multiple comparisons with Bonferroni correction. The chi-square test was used to test the distribution differences of categorical variables. Additionally, we did not adjust for patients’ age because it is strongly associated with tumor types^6^. Root-mean-square error (RMSE) was used to assess deconvolution differences of cell type proportions by groups. Data analysis and visualization were performed using R v.4.2.0. Two-sided *P* < 0.05 was considered statistically significant. *, *P* < 0.05; **, *P* < 0.01; ***, *P* < 0.001; ****, *P* < 0.0001.

